# MicroRNA-24-3p promotes skeletal muscle differentiation and regeneration by regulating HMGA1

**DOI:** 10.1101/2020.11.10.371872

**Authors:** Paromita Dey, Miles A. Soyer, Bijan K. Dey

## Abstract

Numerous studies have established the critical roles of microRNAs in regulating post-transcriptional gene expression in diverse biological processes. Here, we report on the role and mechanism of miR-24-3p in skeletal muscle differentiation and regeneration. miR-24-3p promotes myoblast differentiation and skeletal muscle regeneration by directly targeting high mobility group AT-hook 1 (HMGA1) and regulating it and its direct downstream target, the inhibitor of differentiation 3 (ID3). miR-24-3p knockdown in neonatal mice increases PAX7-positive proliferating muscle stem cells (MuSCs) by derepressing *Hmga1* and *Id3*. Similarly, inhibition of miR-24-3p in the tibialis anterior muscle prevents *Hmga1* and *Id3* downregulation and impairs regeneration. These findings provide evidence that the miR-24-3p/HMGA1/ID3 axis is required for MuSC differentiation and skeletal muscle regeneration *in vivo*.

## Introduction

During postnatal skeletal muscle development and regeneration, quiescent MuSCs are activated to re-enter the cell cycle, followed by proliferation to form a pool of myoblasts, which then differentiate and fuse into newly formed or existing myofibers [1]. This extensive process of making new muscle is known as myogenesis and is essential for maintaining normal physiological function. Dysregulation of the critical molecular and cellular events linked to skeletal myogenesis is associated with muscle degenerative diseases. Most of the understanding of the critical myogenic processes, including myoblast differentiation and skeletal muscle regeneration, is based on the regulation of myogenic transcription factors and signaling molecules [1–8]. However, it is not well understood how these factors of the myogenic network are regulated to maintain normal myogenesis and skeletal muscle function. In this context, an emerging research area is exploring microRNAs’ role in post-transcriptional gene regulation during skeletal myogenesis.

Recently we have begun to elucidate the role of microRNAs in skeletal muscle differentiation, development, and muscle degenerative diseases. The microRNAs involved in myogenesis have recently been reviewed [9]. Most studies have demonstrated the roles of microRNAs in muscle differentiation by overexpression and knockdown experiments using the C2C12 myoblast cell line [10–25]. miR-1 and miR-206, which share identical target binding seed sequences, promote myoblast differentiation by regulating several targets, including *Hdac4*, *Pax7*, *Smarcd2*, and *Smarcb1* [14, 17, 26]. miR-26a promotes myoblast differentiation by regulating the TGFβ/BMP pathway transcription factors SMAD1 and SMAD4 [27]. miR-181 induces muscle differentiation by repressing HOXA11 [28]. In addition, we have demonstrated that several microRNAs, including miR-26a, miR-206, miR-322/424, miR-378, miR-486, and miR-503, are induced during myoblast differentiation and promote myoblast differentiation by regulating critical myogenic regulatory factors and signaling molecules [14, 21–24, 27, 29]. Intriguingly, we have shown that H19 noncoding RNA generates miR-675-3p and miR-675-5p to promote muscle differentiation [29]. Through ChIP-Seq analyses, we have revealed how crucial myogenic transcription factors, including MYOD and MEF2C, repress their repressors by inducing myogenic microRNAs [14, 24], suggesting that microRNAs tightly regulate many critical factors of the myogenic network and maintain the balance between myoblast proliferation and differentiation. However, only a few studies have revealed microRNA function in skeletal muscle development and regeneration in animals [26, 27, 29–36]. Therefore, it is crucial to uncover microRNA function in myogenesis *in vivo* to identify new therapeutic targets for muscle degenerative diseases.

Here, we report that miR-24-3p is abundantly expressed in skeletal muscles and upregulated during myoblast differentiation and muscle regeneration. Consistent with our findings, an earlier study showed that miR-24-3p promotes muscle differentiation *in vitro* [37]. However, it has remained unknown how miR-24-3p promotes myoblast differentiation and whether it plays any role during myogenesis *in vivo*. We show that miR-24-3p promotes myoblast differentiation by directly targeting *Hmga1*and regulating it and its direct downstream target, *Id3*. HMGA1 is a prominent member of the high mobility group A (HMGA) proteins preferentially expressed in the proliferating embryonic tissues but usually absent in differentiated cells [38, 39]. HMGA1 downregulation is crucial for myoblast differentiation [40]. Sustained expression of HMGA1 in myoblast cells represses promyogenic genes, including *Myod* [40]. HMGA1’s direct target ID3 represses MYOD activity by binding and sequestering E12/47 away from the MYOD binding sites of the promyogenic genes, thereby repressing myogenesis. Our findings suggest that miR-24-3p promotes myogenesis by increasing MYOD activity through the downregulation of HMGA1 and ID3. Furthermore, knockdown of miR-24-3p in a neonatal mouse model increases the PAX7-positive proliferating MuSCs by derepressing *Hmga1* and *Id3*. In addition, we show that miR-24-3p is required for skeletal muscle regeneration after injury. Thus, our study establishes a role for miR-24-3p in skeletal muscle function *in vivo* and elucidates the molecular mechanism by which this myogenic microRNA affects muscle differentiation and regeneration: inhibiting a well-established repressor of myogenesis.

## Materials and Methods

### Cell culture

Mouse C2C12 myoblast cell line was purchased from the American Type Culture Collection (ATCC) [41]. These cells were cultured subconfluently in a growth medium (GM) consisting of DMEM (ATCC) supplemented with 10% heat-inactivated fetal bovine serum (FBS) (ATCC) and 1X Antibiotic-Antimycotic (Thermo Scientific). These cells were differentiated into myocytes or myotubes in a differentiation medium (DM) [42]. The DM consisted of DMEM containing 2% heat-inactivated horse serum (Hyclone) and 1X Antibiotic-Antimycotic (Thermo Scientific). Mouse primary myoblast cells were isolated from C57BL/6J mice following a standard protocol described earlier [43]. Mouse primary myoblast cells were differentiated as described [14]. U2OS cells were cultured in DMEM (ATCC) supplemented with 10% heat-inactivated FBS (ATCC) and 1X Antibiotic-Antimycotic (Thermo Scientific). Muscle stem cells (MuSCs) were isolated from mouse TA muscle using a satellite cell isolation kit (Miltenyi Biotec) following the manufacturer’s instructions.

### Isolation of total RNA and performance of qRT–PCR

We isolated total RNAs using Trizol reagent (Invitrogen) or RNeasy mini kit (Qiagen) following the manufacturer’s instructions. We carried out cDNA synthesis using the iScript cDNA Synthesis Kit (Bio-Rad) as instructed. We performed qRT-PCR using the SYBR Green PCR master mix (Bio-Rad) in a Bio-Rad thermal cycler using specific primers for Myogenin (*Myog*) (Forward: AGCGCAGGCTC AAGAAAGTGAATG; Reverse: CTGTAGGCGCTCAATGTACTGGAT), Myosin Heavy Chain (*Mhc*) (Forward: TCCAAACCGTCTCTGCACTGTT; Reverse: AGCGTACAAAGTG TGGGTGTGT), *Hmga1* (Forward: CGACCAAAGGGAAGCAAGAATAA; Reverse: TCCTCTTCC TCCTTCTCCAGTTTC), and *Id3* (Forward: CTCTTAGCCTCTTGGACGACATGA; Reverse: TGT AGTCTATGACACGCTGCAGGA). We used *Rsp13* (Forward: TGACGACGTGAAGGAACAG ATTT; Reverse: ATTTCCAGTCACAAAACGGACCT) or *Gapdh* (Forward: ATGACATCAAGAAG GTGGTGAAGC; Reverse: GAAGAGTGGGAGTTGCTGTTGAAG) primer pairs as a housekeeping gene for normalizing the values of *Myog, Mhc, Hmga1*, and *Id3*. We carried out qRT-PCR for miR-24-3p using the miRCURRY LNA kit as described (Qiagen). The primer sequences used for U6Sn, miR-24-3p, and miR-192 were CTGCGCAAGGATGACACGCA, TGGCTCAGTTCAGCAGGAACAG, and CTGACCTATGAATTGACAGCC, respectively.

### Plasmid construction

Mouse *Hmga1* 3’UTR1 was amplified by PCR from C2C12 myoblast genomic DNA and cloned into pRL-CMV. Specific point mutations in *Hmga1* 3’UTRs (shown in Fig 3A) cloned into pRL-CMV vector were generated using site-directed mutagenesis kit (Stratagene). In addition, *Hmga1* ORF without or with its 3’UTR was subcloned into pMSCV retroviral expression vector as described [14]. We made retroviruses using these constructs in HEK-293T cells by cotransfecting helper plasmids using a standard protocol.

### Luciferase assays

We first transfected U2OS cells with miR-24-3p (UGGCUCAGUUCAGCAGG AACAG), miR-24-3p mutant (UCGCACUGUUCUGCAGGAACAG), or GL2 (UCGAAGUAUUCC GCGUACG), and after 24 hours, we transfected the same cells with Renilla luciferase (rr) plasmids containing 3’ UTRs. pGL3 firefly Photinus pyralis (pp) (Promega) was co-transfected as an internal transfection control. Then we harvested the cells after another 48 hours and performed luciferase assays with a dual-luciferase reporter assay system using a GloMax luminometer (Promega) following our established protocol [14, 27]. We first normalized rr values to the co-transfected pp luciferase values. Then we normalized each rr/pp value in the miR-24-3p-transfected samples with the rr/pp values obtained from GL2-transfected samples.

### Western blotting and antibodies

We harvested the cells, washed them with 1X PBS, lysed them in NP40 lysis buffer (50mMTris-HCl, 150mMNaCl, 0.1% NP-40, 5mM EDTA, 10% glycerol) with protease inhibitors cocktail (Sigma), and sonicated using a Bioruptor (Diagenode). We separated proteins in SDS-PAGE, transferred, and immunoblotted with various antibodies. The antibodies used were anti-HMGA1 (dilution 1:500; Cell Signaling), anti-MYOG (dilution 1:500; Santa Cruz) and anti-MHC (dilution 1:3000; Sigma), and anti-GAPDH (dilution 1:10000; Sigma).

### Intraperitoneal injection of neonatal mice

All animal work was done following our Institutional Animal Care and Use Committee (IACUC) protocol. An antagomir specific to miR-24-3p (Ant-24) and a control GL2 antagomir (Ant-NC) was designed following the procedure, as described previously [44] and were synthesized by Horizon. The sequences for Ant-24 and Ant-NC were mU(*)mG(*)mGmC mUmCmAmGmUmUmCmAmG mCmAmGmGmAmAmCmA(*)mG(*)(3’-Chl) and mU(*)mA(*)mUmCmGmCmGmAmGmUmAmC mGmUmCmGmAmG(*)mG(*)mC(*)mC(*)(3’-Chl), respectively. Here, ‘‘m’’ represents a 2’-O-methyl-modified nucleotide, ‘‘*’’ indicates a phosphorothioate linkage, and ‘‘Chl’’ indicates cholesterol moiety linked to the oligos through a hydroxyprolinol linkage. On postnatal day 3, C57BL/6J pups were injected intraperitoneally (IP). We followed NIH guidelines of sex as biological variables (SABV) and used both male and female pups for our experiments. Ant-24 and Ant-NC were administered twice at a dose of 100 mg/kg body weight in 10–15 μl per injection volume. BrdU was injected 4 hours before harvesting the hind leg skeletal muscles on day 6 for qRT–PCR and fresh-frozen (OCT) samples.

### Skeletal muscle regeneration model and TA muscle injection

We developed an injury-mediated mouse model of skeletal muscle regeneration by intramuscular injection of cardiotoxin (CTX) from Naja nigricollis (EMD Millipore) following our established procedure as described earlier [27, 29]. Briefly, about 10-week-old male C57BL/6J mice were injected on TA muscles with 100ul of 10 μM CTX. This high volume of injection materials, pressure, and post-injection massages spread the injected material throughout the TA muscle compartment. However, we analyzed the middle two-thirds of the TA muscle closest to the injection site. We injected 100 μl (15 μg) of Ant-24 into the TA muscle of one leg and 100 μl (15 μg) of Ant-NC into the contralateral leg 3 days after the CTX injection. Mice were euthanized by CO2 and sacrificed by cervical dislocation before harvesting the muscle samples at different time intervals.

### Immunocytochemistry and immunohistochemistry

We carried out immunocytochemistry following our standard protocol described previously [14, 27]. Briefly, we grew the cells on sterile glass coverslips and fixed them with 2% formaldehyde in PBS for 15 min. Next, we permeabilized the cells with 0.2% Triton X-100 and 1% normal goat serum (NGS) in ice-cold PBS for 5 minutes and blocked them with 1% NGS in PBS twice for 15 minutes. We incubated the fixed and blocked cells with primary antibodies (MYOG 1:50; Santa Cruz Biotechnology and MHC 1:400; Sigma) in 1% NGS for 1 hour. Next, we washed the cells twice with 1X PBS and incubated them with FITC-conjugated anti-mouse IgG (dilution 1:500; Dako Cytomation) for 1 hour. Finally, we rewashed the cells twice and counterstained the nuclei with DAPI while mounting the coverslips on a glass slide (H-1200, Vector Laboratories). Images were captured with an Olympus Hi-Mag microscope. The immunostaining using the skeletal muscle tissue sections was carried out with slight modifications of the protocols described by others: anti-BrdU and LAMININ [45], and anti-DESMIN [34]. The primary antibodies, mouse BrdU (1:50; Roche), rat anti-LAMININ (1:100; Millipore), mouse anti-DESMIN (1:200; Dako), rat anti-F4/80 (1:200; Biolegend), rat anti-CD45 (1:200; Invitrogen), and rat anti-CD4 (1:100; Invitrogen) were used. The secondary antibodies, Alexa Fluor 594 goat anti-mouse IgG1 (1:400; Invitrogen), and Alexa Fluor 488 goat anti-rabbit IgG (1:400; Invitrogen), were used. Images were captured using Zeiss LSM-710 confocal microscope. H&E staining was carried out using a standard protocol, and bright-field images were captured using an Olympus microscope.

### Statistical analyses

Data are presented as the mean+/− standard deviation of three or more biological replicates. A two-tailed Student’s t-test was employed to determine P-values.

## Results

### miR-24-3p is abundantly expressed in adult skeletal muscle and is regulated during myoblast differentiation and skeletal muscle regeneration

An earlier study showed that miR-24-3p induces myoblast differentiation *in vitro* [37]. However, the molecular mechanism of miR-24-3p in myoblast differentiation and its function in skeletal myogenesis *in vivo* remained undetermined. Therefore, we focused on miR-24-3p to reveal its function and molecular mechanism in skeletal myogenesis using primary myoblast cells, neonatal mice, and a mouse model of skeletal muscle regeneration. We first confirmed that miR-24-3p was upregulated during C2C12 myoblast differentiation (Suppl Fig 1). Next, we determined that miR-24-3p was upregulated during mouse primary myoblast differentiation, a physiologically more relevant system for studying myogenesis (Fig 1A). We also examined the expression level of miR-24-3p in various mouse tissues and found that it is abundantly expressed in adult skeletal muscle (Fig 1B). We then examined the expression of miR-24-3p during skeletal muscle degeneration and regeneration. We generated a well-accepted cardiotoxin (CTX)-induced mouse skeletal muscle regeneration model [46] (Fig 1C). We observed a degeneration phase after the CTX injury when quiescent MuSCs are activated and give rise to myoblast cells (Fig 1C). This is followed by a regeneration phase when myoblasts differentiate and fuse to the existing myofibers or make new myofibers (Fig 1C). miR-24-3p decreased rapidly on days 1 through 3 (degeneration phase) and increased on days 5 through 14 (regeneration phase) (Fig 1D). These findings suggest the role of miR-24-3p in myoblast differentiation and skeletal muscle regeneration.

**Figure 1:**
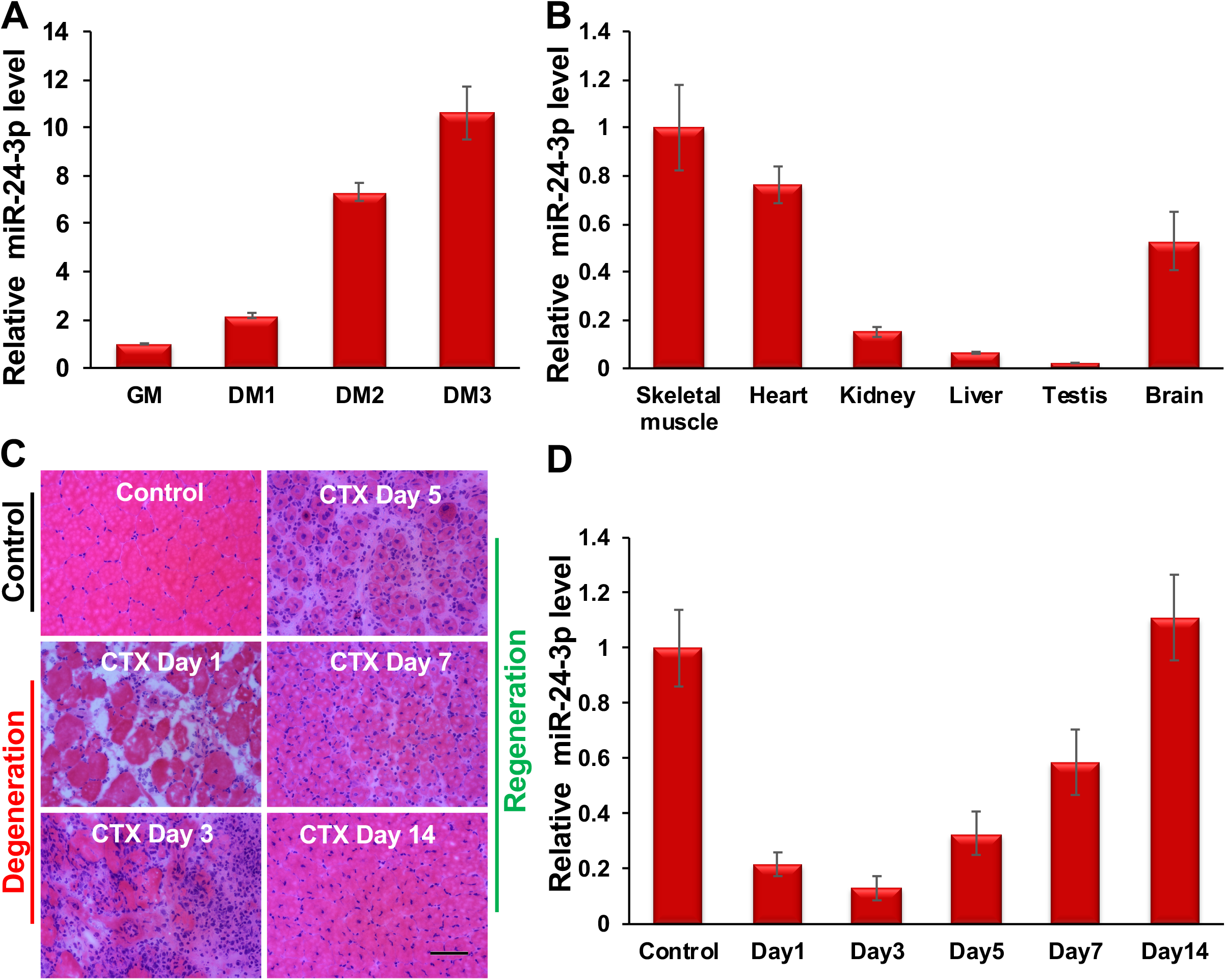
miR-24-3p is abundantly expressed in adult skeletal muscle and upregulated during myoblast differentiation and muscle regeneration. (A) qRT-PCR analyses show that miR-24-3p is upregulated during mouse primary myoblast differentiation. miR-24-3p values were first normalized to the U6sn values, and the fold-change of miR-24-3p was determined relative to the undifferentiated myoblasts (GM). DM1, DM2, and DM3 indicates the number of days myoblasts cultured in differentiation medium (DM). (B) miR-24-3p is abundantly expressed in adult skeletal muscle. qRT-PCR was performed using various mouse tissues to measure miR-24-3p. The values were normalized to the respective U6sn values and again to the values of skeletal muscle. (C) Hematoxylin & eosin (H&E) staining of tibialis anterior (TA) muscle cross-sections from noninjected control and on days 1, 3, 5, 7, and 14 post-injury caused by intramuscular injection of CTX. (D) miR-24-3p is downregulated on days 1–3 post-injury and upregulated on days 5–14 post-injury. The values are expressed as Mean ± SD of three (A-B) and five (D) biological replicates.

### miR-24-3p promotes myoblast differentiation

Since myoblast differentiation is a critical step in postnatal muscle development and regeneration, we assessed the function of miR-24-3p in primary myoblast differentiation. We first confirmed an earlier report that miR-24-3p promotes C2C12 myoblast differentiation (Suppl Fig 2). Next, we examined its function during mouse primary myoblast differentiation. We transfected primary myoblast cells with mature miR-24-3p or GL2, an RNA duplex of the identical length of miR-24-3p from the luciferase gene, as a negative control (NC) in the growth medium (GM). We transfected the cells twice at 24-hour intervals in GM and replaced the GM with the differentiation medium (DM). We carried out immunostaining of the cells for an early myogenic marker, myogenin (MYOG), and a late myogenic marker, myosin heavy chain (MHC), 24 and 48 hours after adding the DM, respectively. The exogenous miR-24-3p increased differentiation, as seen by the increased number of MYOG- and MHC-positive cells with elongated and multinucleated cellular morphology (Fig 2A, B; Suppl Table 1). We also observed increased *Myog* and *Mhc* mRNA and protein levels in these cells when maintained throughout in GM (Fig 2C, D). These results demonstrate that miR-24-3p indeed promotes myoblast differentiation.

**Figure 2:**
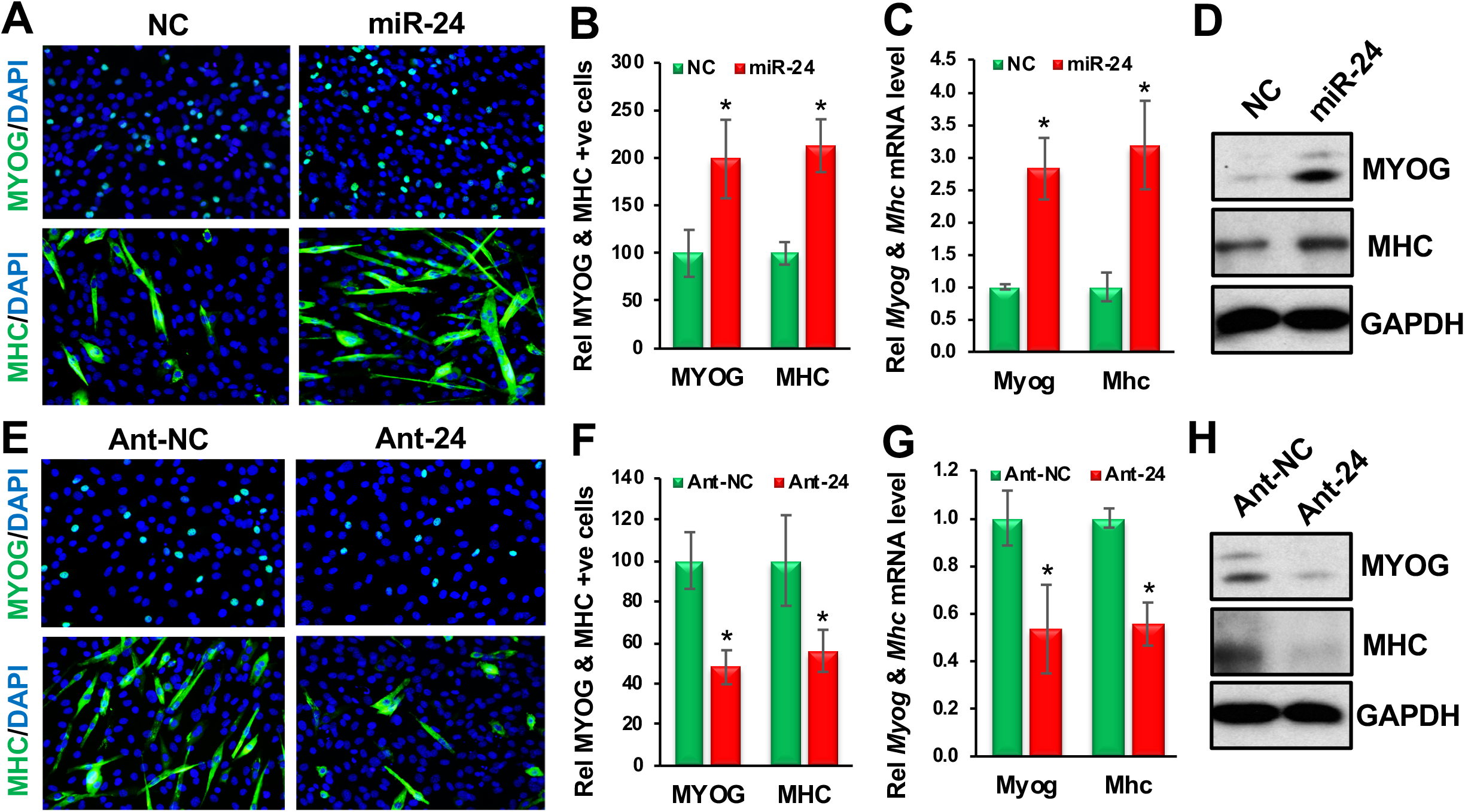
miR-24-3p promotes and is required for primary myoblast differentiation. (A) Primary myoblast cells were transfected twice at 24-hour intervals with GL2 or miR-24-3p in GM, and then the GM was replaced with DM. Immunostaining of these cells was carried out for MYOG and MHC 24 and 48 hours after replacing the GM with DM, respectively. (B) Fractions of MYOG- and MHC-positive cells are presented relative to the GL2 (NC) as 100%. (C) *Myog* and *Mhc* mRNA levels when the cells were maintained throughout in GM. qRT-PCR was carried out for *Myog* and *Mhc*, and the values were normalized to *Gapdh*, then again to the NC values. (D) MYOG, MHC, and GAPDH protein levels. GAPDH served as a loading control. (E) Primary myoblast cells were transfected twice at 24-hour intervals with antagomirs specific to GL2 (Ant-NC) or miR-24-3p (Ant-24), then the GM was replaced with DM. Immunostaining of these cells was carried out for MYOG and MHC 24 and 48 hours after replacing the GM with DM, respectively. (F) Fractions of MYOG- and MHC-positive cells are presented relative to the Ant-NC control as 100%. (G) qRT-PCR was carried out for *Myog* and *Mhc*, and the values were normalized as C. (H) MYOG, MHC, and GAPDH protein levels. GAPDH served as a loading control. The values are expressed as Mean ± SD of biological triplicates. *P < 0.001.

**Figure 3:**
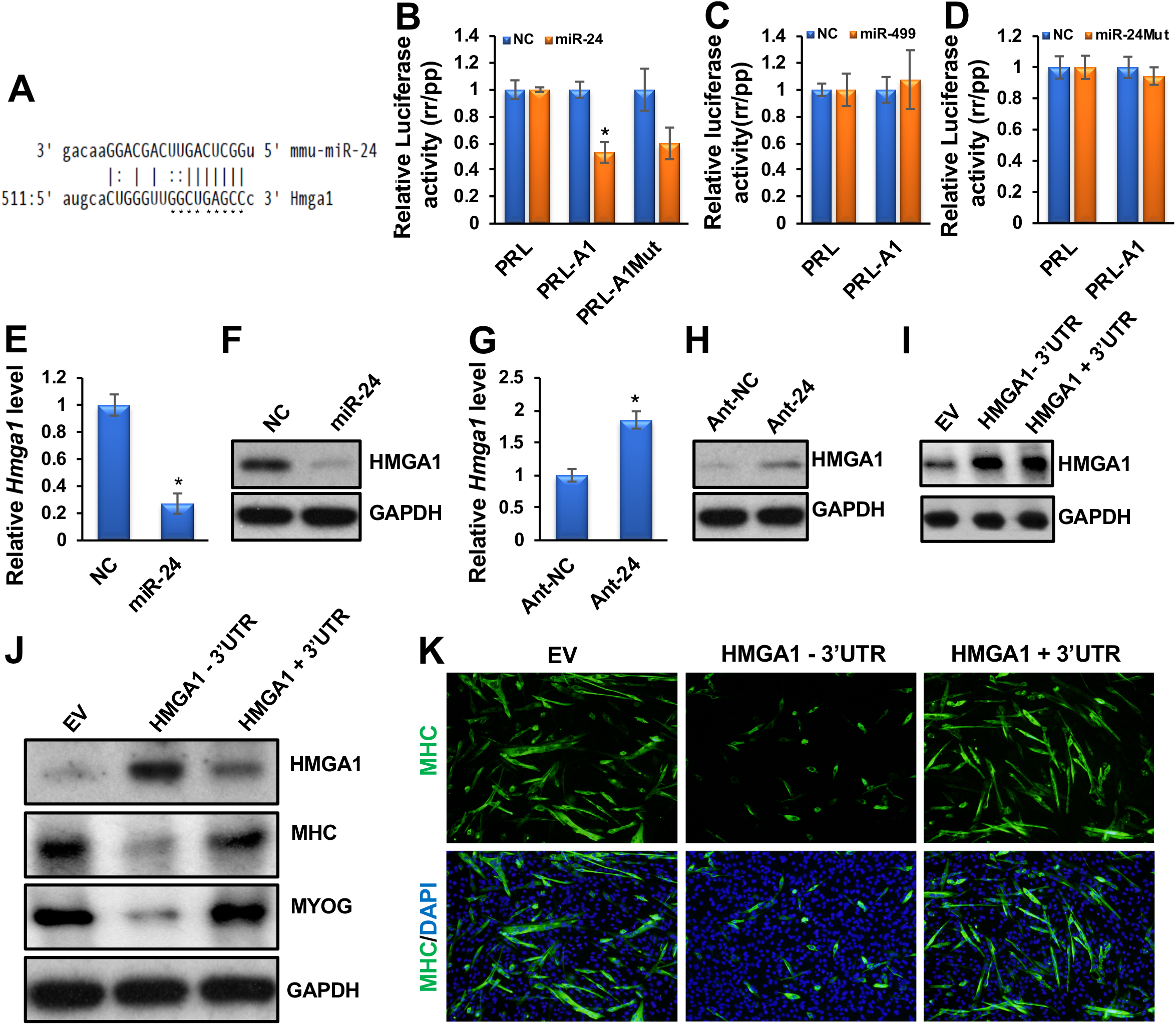
miR-24-3p promotes myoblast differentiation by directly targeting and regulating *Hmga1*. (A) The microRNA target prediction program miRanda predicts a miR-24-3p binding site on the *Hmga1* 3’UTR. The stars indicate the bases that were mutated (PRL-A1Mut). (B) Luciferase assays were performed to measure the effect of miR-24-3p transfection on a Renilla (rr) luciferase reporter fused to the wild-type and mutated *Hmga1* 3’UTRs (PRL-A1 and PRL-A1-Mut), respectively. A firefly (pp) luciferase plasmid was co-transfected as a transfection control. The rr/pp value was normalized with a control rr luciferase plasmid without an *Hmga1* 3’UTR (PRL) and is expressed relative to the normalized rr/pp value in GL2-transfected cells (NC). (C) The luciferase activity was similarly determined when NC or miR-499 was co-transfected with the PRL-A1 construct. (D) An NC or a mutant form of miR-24-3p (miR-24Mut) was co-transfected with PRL or PRL-A1, and luciferase activity was similarly determined. (E, F) Exogenous expression of miR-24-3p downregulated *Hmga1* mRNA and protein levels. (G, H) Ant-24 sustained the expression of *Hmga1* mRNA and protein levels. (I) HMGA1 levels in an empty retroviral vector (EV) or the retroviral vectors stably expressing HMGA1 in myoblasts lacking its 3’UTR (HMGA1-3’UTR) or the 3’UTR attached to it (HMGA1 +3’UTR). (J) Myoblast cells stably expressing EV and the constructs lacking or containing the *Hmga1* 3’UTR were differentiated for 3 days. HMGA1, MYOG, MHC, and GAPDH protein levels are shown. GAPDH served as a loading control. (K) MHC immunostaining on these cells. The values are expressed as Mean ± SD of biological triplicates. *P < 0.001.

We also investigated whether miR-24-3p is required for myoblast differentiation. First, we examined the silencing efficiency of antagomirs specific to miR-24-3p (Ant-24). We transfected primary myoblast cells twice with Ant-24 or control antagomir (Ant-NC) at 24-hour intervals in GM and harvested the cells after another 48 hours. We determined by qRT-PCR analyses that the endogenous miR-24-3p level was downregulated five-fold in the Ant-24-transfected samples (Suppl Fig 3). Next, we transfected the cells twice with Ant-24 or Ant-NC at 24-hour intervals in GM and replaced the GM with the DM. We carried out immunostaining of the cells for MYOG and MHC 24 and 48 hours after adding the DM, respectively. Ant-24 decreased the MYOG- and MHC-positive cell numbers and differentiation morphology (Fig 2E, F; Suppl Table 1). We also found that both *Myog* and *Mhc* mRNA and protein levels were significantly decreased in these samples (Fig 2G, H). We followed up these cells for five days (Suppl Fig 4). Whereas Ant-NC-treated cells differentiate normally and form multinucleated myotubes (Suppl Fig 4A, D), the Ant-24-treated cells don’t differentiate following the normal kinetics (Suppl Fig 4B, E) and the majority of the cells looks like proliferating myoblasts (Suppl Fig 4C, F). These findings establish that miR-24-3p is required for myoblast differentiation.

### miR-24-3p promotes muscle differentiation by directly targeting and regulating Hmga1

To determine how miR-24-3p promotes myoblast differentiation, we revealed its direct target relevant to myogenesis. The microRNA target prediction algorithm miRanda revealed *Hmga1* as a potential target for miR-24-3p (Fig 3A). HMGA1 is a known myogenic repressor, and its downregulation is required for myoblast differentiation [40]. Therefore, we chose *Hmga1* to investigate whether miR-24-3p promotes muscle differentiation by directly targeting and regulating *Hmga1*. To determine if *Hmga1* 3’UTR has a valid target site(s) for miR-24-3p, we fused the *Hmga1* 3’UTR to a luciferase reporter gene driven by the cytomegalovirus (CMV) promoter. Transfection of miR-24-3p with the *Hmga1* 3’UTR luciferase construct repressed luciferase activity (Fig 3B). However, mutations in the seed sequence of the predicted target site in the *Hmga1* 3’UTR did not relieve the repression significantly (Fig 3B), suggesting a non-canonical (non-seed-match) target site(s) for miR-24-3p in the *Hmga1* 3’UTR. Non-seed-match target sites have previously been reported by us and others [14, 47]. Consistent with this notion, the RNA hybrid target prediction program revealed several non-canonical targets sites of miR-24-3p in the *Hmga1* 3’UTR (Suppl Fig 5).

In addition to the conventional experiments as described above, we carried out additional control experiments where we co-transfected *Hmga1* 3’UTR with a control miR-499 (similarly upregulated during myoblast differentiation but does not contain *Hmga1* target sites) and miR-24Mut (randomly mutated miR-24-3p at the seed nucleotide positions 2, 5, and 7 and a non-seed nucleotide position 12). We changed one base outside the seed sequence as non-seed sequences also influence microRNA binding to the target sites. Co-transfection of the 3’UTR of *Hmga1* either with miR-499 or mutated form of miR-24-3p did not suppress the luciferase activity (Fig 3C, D), further suggesting that *Hmga1* is specifically targeted by miR-24-3p.

To demonstrate that *Hmga1* is indeed a valid target of miR-24-3p during myogenesis, we transfected primary myoblasts with GL2 or miR-24-3p twice at 24-hour intervals in GM. We harvested the cells for qRT-PCR and Western blotting for HMGA1 48 hours after the last transfection. miR-24-3p drastically downregulated *Hmga1* mRNA and protein levels in these samples (Fig 3E, F). We also observed downregulation of HMGA1’s direct target *Id3* transcripts in these samples (Suppl Fig 6A). HMGA1 is known to bind to the *Id3* promoter and regulate its expression positively [48]. Consistent with this finding, knockdown of *Hmga1* in myoblasts decreased the *Id3* level (Suppl Fig 6B, C). In a reciprocal experiment, inhibiting miR-24-3p in DM caused derepression of endogenous *Hmga1* mRNA and protein (Fig 3G, H) and *Id3* transcript (Suppl Fig 6D) levels during primary myoblast differentiation. These findings demonstrate that miR-24-3p is responsible for the repression of *Hmga1* mRNA and protein levels during myoblast differentiation.

We further examined if the 3’UTR of *Hmga1* containing the miR-24-3p target site was necessary for downregulation of the *Hmga1* mRNA and protein levels during myoblast differentiation. We used retroviral vectors stably expressing *Hmga1* lacking its 3’UTR (HMGA1-3’UTR) or the 3’UTR attached to it (HMGA1+3’UTR) during myoblast differentiation. We also used myoblasts stably expressing empty vectors (EV). We chose stable cells containing either HMGA1-3’UTR or HMGA1+3’UTR constructs expressing a similar basal level of HMGA1 for differentiation assays (Fig 3I). After three days in DM, HMGA1 protein persisted in the cells expressing *Hmga1* without its 3’UTR, in contrast to the cells expressing *Hmga1* containing the 3’UTRs or an EV (Fig 3J). As expected, the myoblast cells expressing *Hmga1* attached to its 3’UTR or an EV normally differentiate, as found by immunoblotting for MYOG and MHC and immunofluorescence for MHC (Fig 3J, K). However, the cells expressing *Hmga1* without its 3’UTR repress differentiation (Fig 3J, K). Together, these findings confirm that *Hmga1* is a bonafide target for miR-24-3p and that miR-24-3p promotes myoblast differentiation by directly targeting and regulating *Hmga1*.

### miR-24-3p regulates proliferation of PAX7-positive MuSCs in vivo

To determine the role of miR-24-3p *in vivo*, we knocked down endogenous miR-24-3p in the skeletal muscle of neonatal mice using antagomirs [44]. We injected 3-day-old neonatal mice intraperitoneally with antagomirs specific to miR-24-3p (Ant-24) or NC (Ant-NC) twice at 24-hour intervals and harvested the muscles on day 6. Ant-24 knocked down miR-24-3p in the hind leg skeletal muscles but had no impact on control miR-192 (Fig 4A). The *Hmga1* mRNA level was derepressed in the miR-24-3p knockdown skeletal muscles (Fig 4B). HMGA1 activity was upregulated in these muscles, as indicated by an increase in *Id3* transcripts (Fig 4B). Ant-24 decreased the endogenous miR-24-3p level in the MuSCs isolated from these samples (Fig 4C). Consequently, *Myog* and *Mhc* levels were decreased in the undifferentiated and differentiated MuSCs isolated from these samples (Suppl Fig 7A, B). To examine the effects of miR-24-3p knockdown on MuSC proliferation, we injected BrdU into these mice to label the proliferating cells 4 hours before harvesting the muscles. We observed a significant increase in BrdU-positive cells in Ant-24 skeletal muscles (Fig 4D, E). The BrdU-positive cells were located at the periphery of the muscle fibers, and 90.87% of these cells also expressed the MuSC marker PAX7 in Ant-24 samples (Fig 4F). Based on these results, we conclude that miR-24-3p restricts the proliferative potential of MuSCs and induces differentiation *in vivo*. Thus, our findings suggest that the miR-24-3p/HMGA1/ID3 axis plays an essential role during neonatal skeletal muscle development by regulating MuSC proliferation.

**Figure 4:**
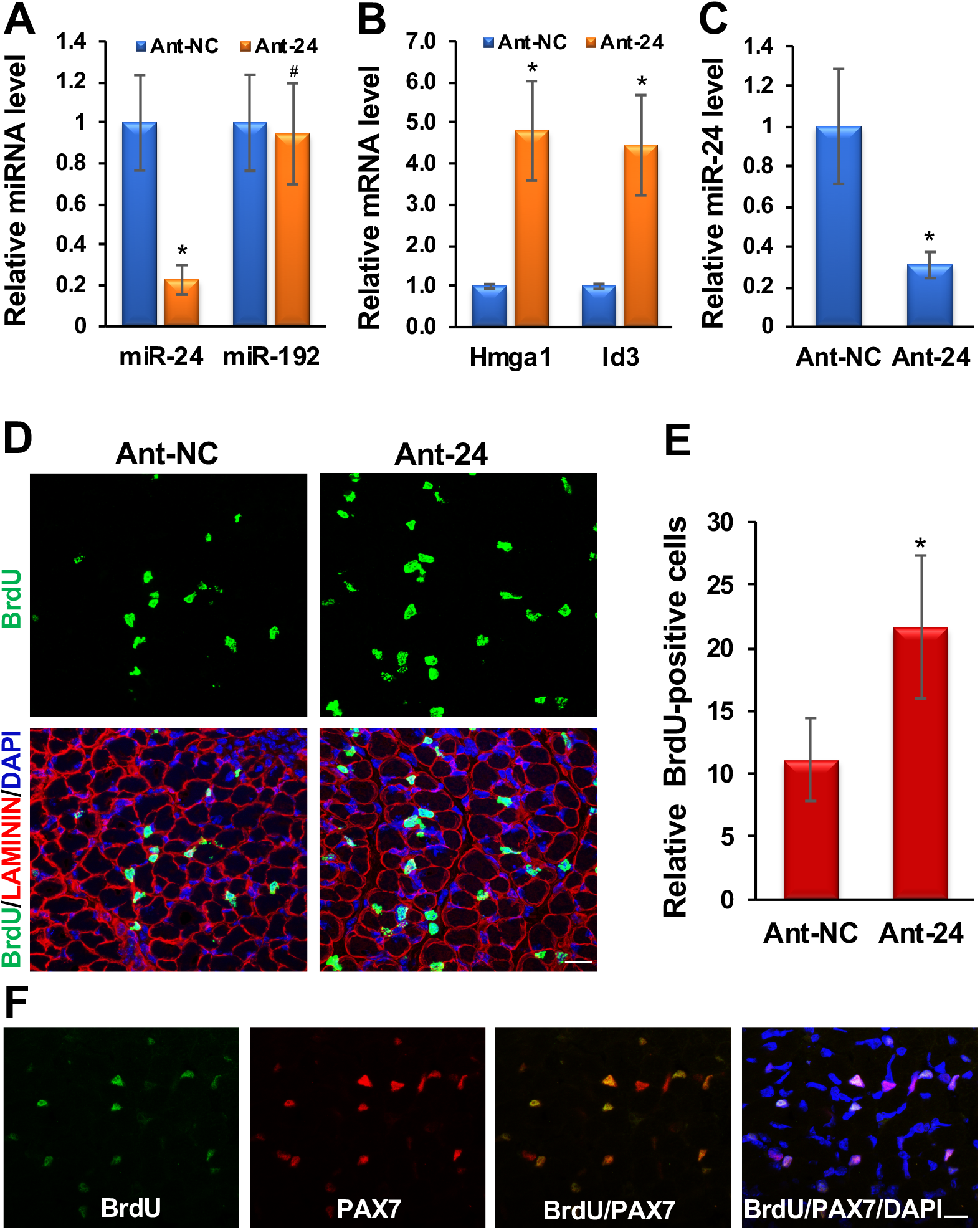
Knockdown of miR-24-3p increases PAX7-positive proliferating MuSCs *in vivo*. (A) Intraperitoneal injection of Ant-24 decreases endogenous miR-24-3p in neonatal skeletal muscle. qRT-PCR values of miR-24-3p normalized to the U6sn values are expressed relative to Ant-NC values. miR-192 was used as an NC. (B) Ant-24 derepresses *Hmga1* and *Id3* transcript levels. qRT-PCR of *Hmga1* and *Id3* on hind leg skeletal muscle normalized to *Gapdh*, then again to the respective Ant-NC samples. (C) miR-24-3p level is decreased in the MuSCs isolated from Ant-24-injected neonatal mice. (D) Confocal images of skeletal muscle harvested 4 hours after BrdU labeling from Ant-24- or Ant-NC-injected neonates. Cell proliferation was determined by anti-BrdU antibody (green), the cell surface was marked by LAMININ (red), and nuclei were counterstained with DAPI (blue). (E) Relative BrdU-positive nuclei per field. Mean ± SD of 10 random fields from five mice. (F) Confocal microscopy images of Ant-24-injected neonatal skeletal muscles from D immunostained for BrdU (green), PAX7 (red), DAPI (blue), BrdU and PAX7 (orange), and BrdU, Pax7, and DAPI (magenta). Mean ± SD of the samples from five neonatal mice. *P < 0.001. Scale Bar: 50 μM.

### miR-24-3p is essential for adult skeletal muscle regeneration

We used a well-established mouse model of skeletal muscle regeneration by intramuscular injection of CTX into the tibialis anterior (TA) muscle. This model first induces injury (degeneration) and later regenerates spontaneously, allowing us to study muscle regeneration (Fig 1C). As described earlier, miR-24-3p decreased rapidly on days 1–3 after CTX injury, then gradually increased as new muscles were formed (Fig 1D). We examined the *Hmga1* levels in these samples and found its expression pattern anti-correlated to miR-24-3p, increasing on days 1–3 after injury and decreasing steadily on days 5–14 after injury (Fig 5A). These findings suggest that miR-24-3p regulates *Hmga1* during regeneration.

**Figure 5:**
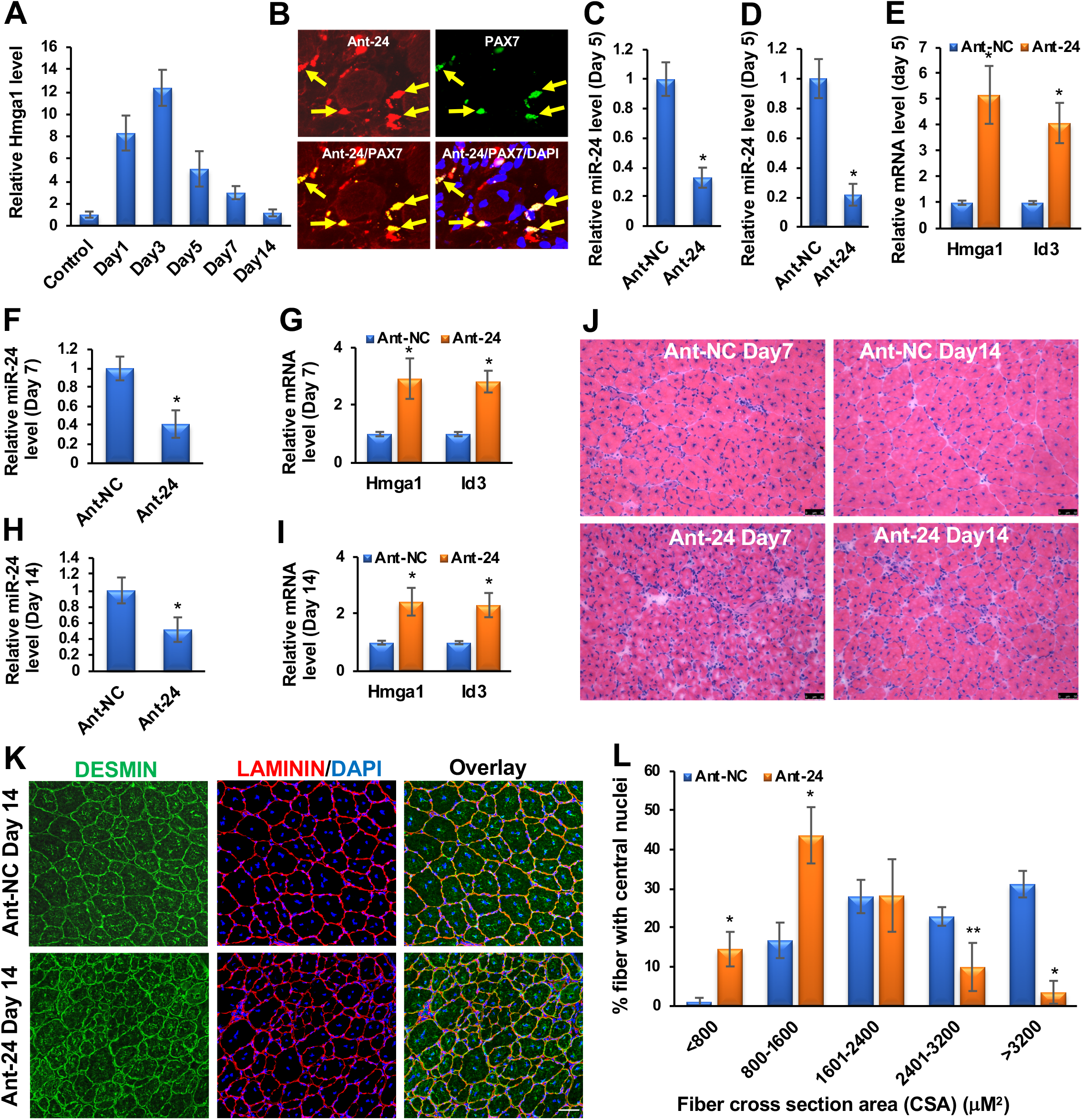
miR-24-3p is essential for skeletal muscle regeneration. (A) *Hmga1* is gradually upregulated in TA muscles on days 1–3 post-injury and downregulated on days 5–14 post-injury. *Hmga1* values were normalized to the *Rps13* values. (B) We injected a fluorescence (DyLight594) conjugated Ant-24 into TA muscle of adult mice on day 3 after CTX injury and harvested the samples day 5 post-injury. Ant-24 incorporates into TA muscle, including MuSCs on day 5 postinjury. Arrows indicate a subset of PAX7-positive MuSCs overlapped with Ant-24 fluorescence. (C) miR-24-3p level was also decreased in the MuSCs isolated from these samples. qRT-PCR was carried out for miR-24-3p, and its values were first normalized to the U6sn values and again relative to NC values. (D, E) Ant-NC and Ant-24 were injected in TA muscle on day 3 post-injury. The miR-24-3p level was repressed, and *Hmga1* and *Id3* transcript levels were derepressed in the TA muscle on day 5 post-injury. *Hmga1* and *Id3* values were normalized to the respective *Rps13* values and then again to the Ant-NC values. (F-I) the miR-24-3p level remained repressed on days 7 and 14 post-injury, and *Hmga1* and *Id3* transcript levels remained deprepressed in the Ant-24 samples. (J) Representative images of H&E staining showing that the Ant-24 impairs regeneration on days 7 and 14 post-injury. (K, L) DESMIN and LAMININ staining and crosssection areas (CSA) of the regenerating fibers on day 14 using the Fiji software. More than 600 fibers were counted from ten random sections of five mice in both groups. Scale Bar: 50 μM. Mean ± SD of the samples from five mice. *P < 0.001; **P < 0.01.

Next, we confirmed the role of miR-24-3p during skeletal muscle regeneration by knocking it down using Ant-24. We first determined whether Ant-24 incorporated into MuSCs after injury and decreased endogenous miR-24-3p level by injecting a fluorescence conjugated Ant-24 into TA muscle of adult mice on day 3 post-injury. Ant-24 incorporates into TA muscle, including MuSCs on day 5 post-injury (Fig 5B), and indeed decreased miR-24-3p level in these samples (Fig 5C, D). Furthermore, decreased miR-24-3p level in TA muscle was associated with derepression of *Hmga1* and *Id3* transcript levels (Fig 5E).

Since miR-24-3p was upregulated on days 5–14 after CTX injury, we injected Ant-24 into the TA muscles of one leg and Ant-NC into the TA muscles of the contralateral leg on day 3 post-injury. On day 5 post-injury, the regeneration process was early, when new myofibers were just beginning to form. Therefore, we examined the impact of Ant-24 on days 7 and 14 after injury. The miR-24-3p level remained repressed while the *Hmga1* and *Id3* transcript levels remained derepressed in the Ant-24 samples (Fig 5F-I). In Ant-NC mice, muscle regenerated normally, and the morphology was restored on day 14, except for the presence of the central nuclei, a signature of regenerating muscles (Fig 5J). In contrast, the regeneration process was impaired in the Ant-24 muscles. In addition, we observed the presence of numerous nuclei, indicating increased inflammatory cells among other cell types [49–51] (Suppl Fig 8A-L), and the persistence of smaller myofibers, indicating impaired regeneration (Fig 5J). We further characterized the regenerating skeletal muscle sections on day 14 after injury by DESMIN and LAMININ immunostaining and measuring myofiber cross-section areas (CSA). DESMIN is an intermediate filament protein abundantly expressed in newly generated muscle fibers, while LAMININ labels the muscle fiber boundaries. DESMIN was expressed in the myofibers of regenerating muscles in both Ant-24 and Ant-NC samples (Fig 5K; Suppl Fig 9). However, the muscle fibers were significantly smaller in the Ant-24 muscles than in the Ant-NC muscles (Fig 5L). In addition, interfiber spaces were larger and filled with nuclei, further supporting abnormal regeneration in the Ant-24 muscles. These findings suggest that the miR-24-3p/HMGA1/ID3 axis plays a critical role in normal skeletal muscle regeneration *in vivo*.

## Discussion

Most of the understanding of skeletal muscle development and regeneration is based on the regulation of myogenic transcription factors and signaling molecules [1–8]. However, it remains elusive how these critical factors of the fundamental myogenic processes are themselves regulated. Furthermore, molecular mechanisms underlying the regulation of gene expression during skeletal muscle development and regeneration are not entirely understood, hindering the development of therapeutic interventions for muscle degenerative diseases. To fill this knowledge gap, we have examined the role of a myogenic microRNA in myoblast differentiation and skeletal muscle regeneration. Thus far, most of the studies have demonstrated the role of microRNAs in C2C12 myoblast differentiation *in vitro* [10–25]. However, only a limited number of studies, including ours, have revealed the function of microRNAs using animal models of muscle development and regeneration [26, 27, 29, 34–36]. In this study, we demonstrated the role of miR-24-3p in muscle differentiation and regeneration in mice. An earlier study also suggests that this microRNA is upregulated during myoblast differentiation and promotes differentiation *in vitro* [37]. However, it remains unknown how miR-24-3p promotes myoblast differentiation and whether it has any role in myogenesis *in vivo*. Here we demonstrated that miR-24-3p promotes myoblast differentiation by directly targeting and regulating HMGA1, a well-known repressor of myogenesis [40]. More importantly, we have shown that miR-24-3p is essential for skeletal muscle regeneration. Our study establishes the role of a myogenic microRNA in skeletal muscle function *in vivo*. Also, it provides mechanistic insights into how a myogenic microRNA sculpts myogenesis by regulating a well-established repressor of myogenesis.

HMGA1 is an important member of the high mobility group A (HMGA) proteins predominantly expressed in the proliferating embryonic tissues but mostly absent in differentiated cells [38, 39]. HMGA1 downregulation is essential for myogenic differentiation [40]. Sustained expression of HMGA1 in myoblast cells represses promyogenic genes, including *Myod* [40]. HMGA1’s direct target ID3 represses MYOD activity by binding and sequestering E12/47 away from the MYOD binding sites of the critical promyogenic genes, thereby repressing myogenesis. Therefore, our findings suggest that miR-24-3p promotes myogenesis by increasing MYOD activity through the downregulation of HMGA1 and ID3. Consistent with this idea, HMGA1 is involved in the differentiation of several cell types, including embryonic lymphohematopoietic cells and adipocytes [52, 53]. These findings indicate that the strictly regulated expression of HMGA1 is critical for normal cellular differentiation, including myogenic differentiation. Our study reveals how a myogenic microRNA regulates this crucial factor during myoblast differentiation. Our studies suggest that miR-24-3p/HMGA1/ID3 axis is essential for normal myogenesis. Since the loss of *Hmga1* gene function affects cellular differentiation processes [39] and *Hmga1* knockout mice develop diseases including type 2 diabetes [54], cardiac hypertrophy [55], and myelolymphoproliferative disorders [52], we envisage that dysregulation of miR-24-3p/HMGA1/ID3 axis may lead to muscle degenerative diseases.

Only a handful of studies have implicated microRNAs in skeletal muscle function in animal development [26, 27, 29–36]. This gap prompted us to study the function of miR-24-3p in skeletal muscle differentiation and regeneration *in vivo* using mouse models. Though defective skeletal muscle development in the muscle-specific deletion of Dicer suggests a critical role for microRNAs during skeletal muscle development in animals [30], there have been conflicting findings in the literature on whether specific microRNAs are essential for these processes. In one study, the germline deletion of miR-206 did not result in any significant defects in skeletal muscle development and exerted only a mild effect on skeletal muscle innervation following injury [31]. Although miR-1 is expressed in skeletal muscles, miR-1 knockout mice showed defects only in cardiac muscle development; no visible phenotypes were observed in skeletal muscles [56]. These discrepancies might be due to the functional redundancy between miR-1 and miR-206, as they share identical seed sequences. In another study, miR-206 knockout delayed skeletal muscle regeneration following CTX injury [34]. We demonstrated earlier that miR-26a and H19-derived miR-675-3p and miR-675-5p play essential roles in skeletal muscle development and regeneration [27, 29]. Here, we have demonstrated a critical role of miR-24-3p in myogenesis *in vivo*. It’s important to elucidate the role of the repertoire of microRNAs in myogenesis *in vivo* because their concerted actions maintain the normal skeletal muscle function. Future studies will explore its role in muscle degenerative diseases. Nonetheless, our findings elucidate an essential function of miR-24-3p in skeletal muscle differentiation and regeneration *in vivo*.

## Declarations

### Ethics approval and consent to participate

The animal work of this manuscript was approved by the institutional animal care and use committee (IACUC).

### Consent for publication

The authors give consent for publication in CMLS.

### Availability of data and material

Data generated for this study are included in the manuscript and available from the corresponding author.

### Competing interests

The authors declare no competing interests.

### Funding

This work was supported by SUNY startup and American Heart Association (AHA 17SDG33670339) grants to BKD.

### Authors’ contributions

BKD conceived the project; PD and BKD designed the study; PD, MAS, and BKD performed the experiments; PD and BKD analyzed the data and prepared the manuscript.

## Acknowledgments

The authors acknowledge the inputs of the Dey lab members. We acknowledge Dr. Melinda Larsen for providing F4/80 and CD45 antibodies.

## Notes

### Competing Interest Statement

The authors have declared no competing interest.

### Summary of Updates

We have added additional experimental data.

