## supplemental materials for "MicroRNA-24-3p promotes skeletal muscle differentiation and regeneration by regulating HMGA1"

**Running title:** miR-24-3p regulates skeletal muscle differentiation and regeneration

**Keywords: Keywords:** skeletal muscle stem cell, myoblast, differentiation, development, regeneration, microRNA, miR-24-3p, HMGA1, Id3

The supplementary pdf file includes:

Suppl Table 1: miR-24-3p increases and Ant-24 decreases number of MYOG- and MHC-positive cells.

Suppl Table 2: List of oligos used in this study.

Suppl Fig 1: miR-24-3p is upregulated during C2C12 myoblast differentiation.

Suppl Fig 2: miR-24-3p promotes C2C12 myoblast differentiation.

Suppl Fig 3: miR-24-3p level is downregulated in the Ant-24-transfected primary myoblasts.

Suppl Fig 4: Ant-24 represses primary myoblast differentiation.

Suppl Fig 5: RNA hybrid microRNA target prediction algorithm identifies non-seed match miR-24-3p target sites in the 3'UTR of *Hmga1*.

Suppl Fig 6: miR-24-3p regulates HMGA1's direct downstream target *Id3* transcript level during myoblast differentiation.

Suppl Fig 7: *Myog* and *Mhc* transcript levels are downregulated in the undifferentiated and differentiated MuSCs isolated from Ant-24-injected neonatal mice.

Suppl Fig 8: Increased presence of various cells, including inflammatory cells are observed in the Ant-24-injected regenerating skeletal muscle.

Suppl Fig 9: Ant-24 impairs skeletal muscle regeneration after injury.

| Transfection | MYOG-positive cell number | | | MHC-positive cell number | | |
|---|---|---|---|---|---|---|
| | Total counts (10 fields) | Percentage (%) | | Total counts (10 fields) | Percentage (%) | |
| | | Mean | SD | | Mean | SD |
| NC | 291 | 100 | 24.65 | 256 | 100 | 11.82 |
| miR-24 | 554 | 197.78 | 41.19 | 542 | 213.83 | 27.72 |
| Ant-NC | 259 | 100 | 13.78 | 330 | 100 | 22.04 |
| Ant-24 | 123 | 48.22 | 8.28 | 179 | 55.38 | 10.52 |

**Suppl Table 1: miR-24-3p increases and Ant-24 decreases number of MYOG- and MHC-positive cells.** Primary myoblast cells were transfected with mature miR-24-3p or GL2 as an NC twice at 24-hour intervals in GM, then replaced the GM with DM. Immunostaining on these cells was carried out for an early myogenic marker, myogenin (MYOG), and a late myogenic marker, myosin heavy chain (MHC) 24 and 48 hours after adding the DM, respectively. In a reciprocal experiment, primary myoblast cells were transfected twice with antagomirs specific to miR-24-3p (Ant-24) or GL2 (Ant-NC) at 24-hour intervals in GM, then replaced the GM with DM. Immunostaining on these cells was carried out for MYOG and MHC 24 and 48 hours after adding the DM, respectively. Fractions of MYOG- and MHC-positive cells were determined from 10 random fields, and data are presented relative to the NC or Ant-NC control as 100%.

| Oligos | Sequence |
|---|---|
| U6sn primer | CTGCGCAAGGATGACACGCA |
| miR-24 primer | TGGCTCAGTTCAGCAGGAACAG |
| miR-192 primer | CTGACCTATGAATTGACAGCC |
| Gapdh Forward primer | ATGACATCAAGAAGGTGGTGAAGC |
| Gapdh Reverse primer | GAAGAGTGGGAGTTGCTGTTGAAG |
| Rps13 Forward primer | TGACGACGTGAAGGAACAGATTT |
| Rps13 Reverse primer | ATTTCCAGTCACAAAACGGACCT |
| Myog Forward primer | AGCGCAGGCTCAAGAAAGTGAATG |
| Myog Reverse primer | CTGTAGGCGCTCAATGTACTGGAT |
| Mhc Forward primer | TCCAAACCGTCTCTGCACTGTT |
| Mhc Reverse primer | AGCGTACAAAGTGTGGGTGTGT |
| Hmga1 Forward primer | CGACCAAAGGGAAGCAAGAATAA |
| Hmga1 Reverse primer | TCCTCTTCCTCCTTCTCCAGTTTC |
| Id3 Forward primer | CTCTTAGCCTCTTGGACGACATGA |
| Id3 Reverse primer | TGTAGTCTATGACACGCTGCAGGA |
| miR-24 | UGGCUCAGUUCAGCAGGAACAG |
| miR-24Mut | UCGCACUGUUCUGCAGGAACAG |
| GL2 | UCGAAGUAUUCCGCGUACG |
| Ant-24 | mU(*)mG(*)mGmCmUmCmAmGmUmUmCmAmGmCmAmGmGmA mAmCmA(*)mG(*)(3'-Chl) |
| Ant-NC | mU(*)mA(*)mUmCmGmCmGmAmGmUmAmCmGmUmCmGmAmG (*)mG(*)mC(*)mC(*)(3'-Chl) |

**Suppl Table 2: List of oligos used in this study.**

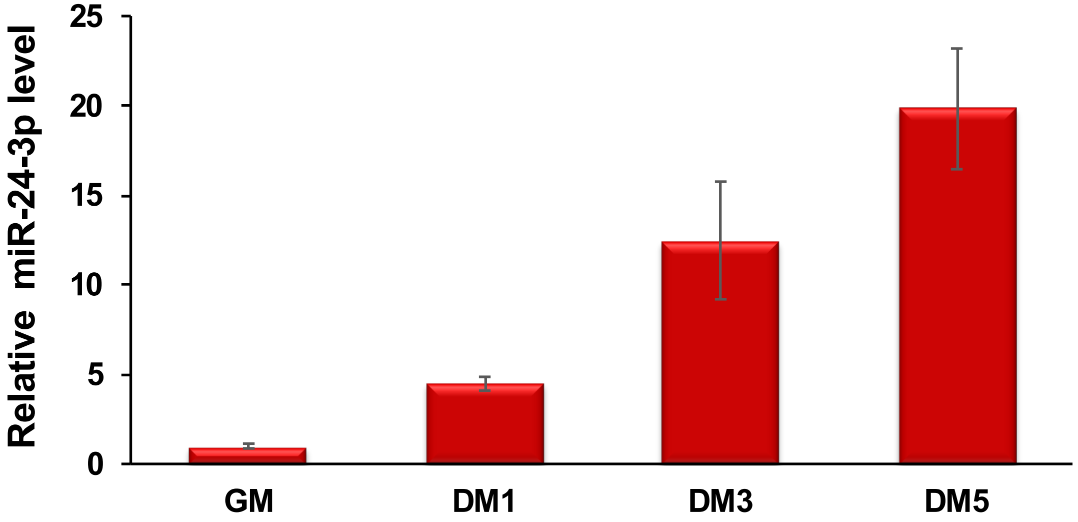


**Suppl Fig 1: miR-24-3p is upregulated during C2C12 myoblast differentiation.** qRT-PCR analyses show that miR-24-3p is upregulated during C2C12 myoblast differentiation. miR-24-3p values were first normalized to the U6sn values, and fold-change of miR-24-3p was determined relative to the undifferentiated myoblasts (GM). DM1, DM3, and DM5 indicates the number of days C2C12 myoblasts cultured in differentiation medium (DM). The values are expressed as Mean ± SD of biological triplicates.

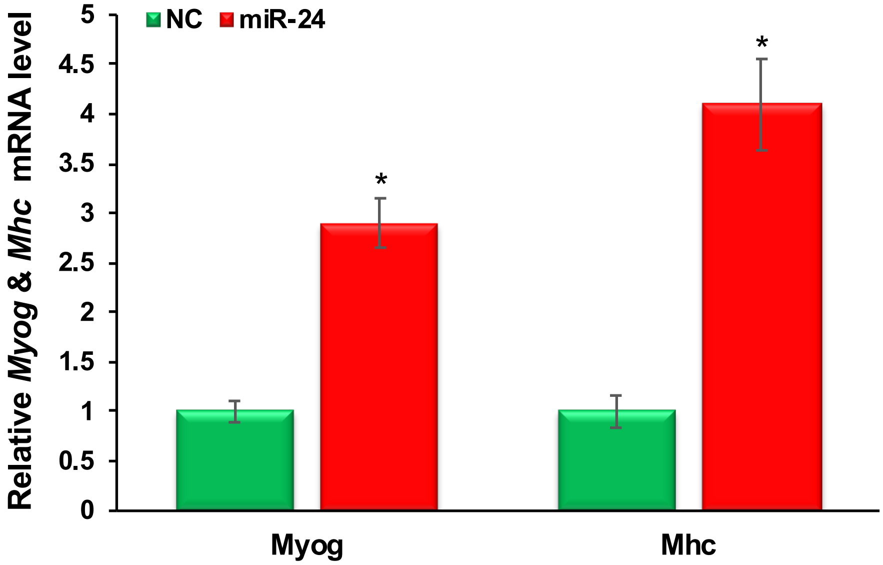


**Suppl Fig 2: miR-24-3p promotes C2C12 myoblast differentiation.** C2C12 myoblast cells were transfected twice at 24-hour intervals with GL2 negative control (NC) or miR-24-3p in GM. The cells were harvested after another 48 hours. qRT-PCR was carried out for *Myog* and *Mhc*, and the values were normalized to *Gapdh*, then again to the values of the NC samples. The values are expressed as Mean ± SD of biological triplicates. *P < 0.001.

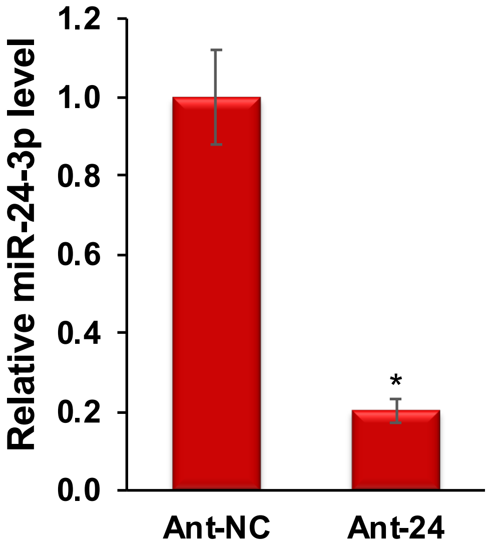


**Suppl Fig 3: miR-24-3p level is downregulated in the Ant-24-transfected primary myoblasts.** We transfected primary myoblast cells twice with antagomirs specific to miR-24-3p (Ant-24) or GL2 control (Ant-NC) at 24-hour intervals in GM and harvested the samples after another 48 hours. We performed qRT-PCR analyses for miR-24-3p. miR-24-3p values were normalized to the respective U6sn values. The values are expressed as Mean ± SD of biological triplicates. *$P < 0.001$.

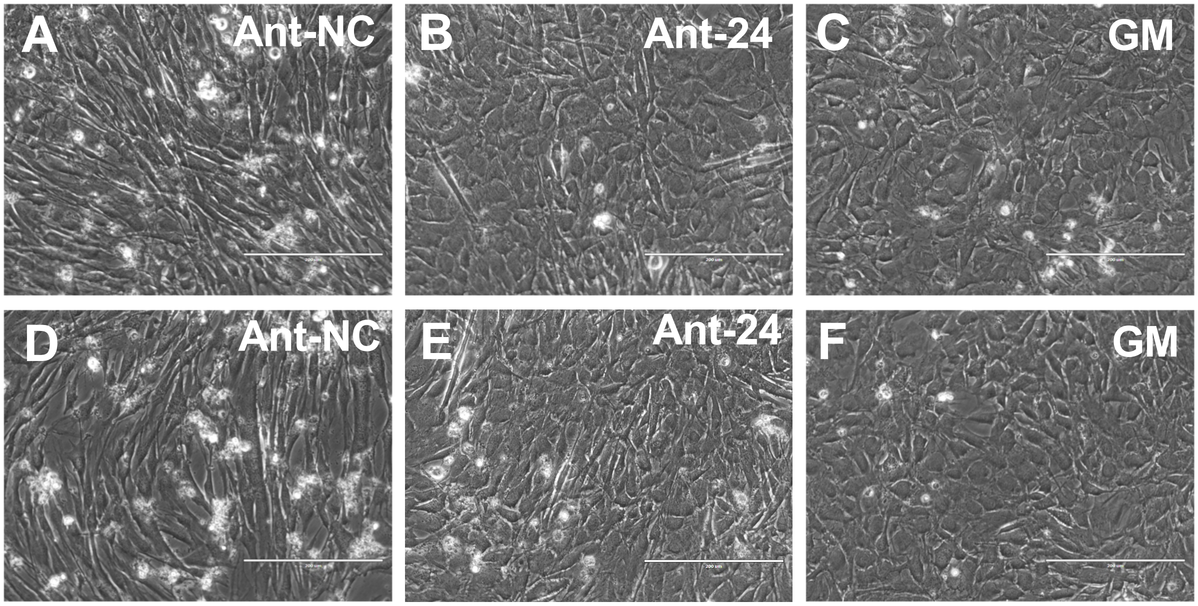


**Suppl Fig 4: Ant-24 represses primary myoblast differentiation.** We transfected primary myoblast cells twice with antagomirs specific to miR-24-3p (Ant-24) or GL2 control (Ant-NC) at 24-hour intervals in GM and replaced the GM with DM. We followed up these cells for 5 days. (A, D) Ant-NC-treated cells differentiate normally and forms multinucleated myotubes. (B, E) Ant-24-treated cells do not differentiate following the normal kinetics, and the majority of the cells looks like (C, F) proliferating myoblasts grown in GM. Scale Bar: 200 μM.

```
position  311
target 5' U   C        UUC       U 3'
           CUG  CCUGCUG   CUG GCU
           GAC  GGACGAC   GAC CGG
miRNA  3'     AA       UU    U   U 5'
```

```
position  182
target 5' C    GCCC    C C    ACACCU   C 3'
           CUGU    UCCU C GAAC      GCC
           GACA    AGGA G CUUG      CGG
miRNA  3'              C A    ACU      U 5'
```

```
position  1087
target 5' U      GUGGG GA   G       G 3'
           UGUUCU     G  GGG UG GCCA
           ACAAGG     C  CUU AC CGGU
miRNA  3' G      A     GA   G  U      5'
```

```
position  500
target 5'    C      CCCACCCU  A   A   U 3'
              UCCUGC        AC GAG UCA
              AGGACG        UG CUC GGU
miRNA  3' GACA      ACU       A         5'
```

```
position  897
target 5' A   UCA    C       U    U 3'
           UGU   UCCU CUGAAUU  GCC
           ACA   AGGA GACUUGA  CGG
miRNA  3' G          C       CU   U 5'
```

```
position  78
target 5'     N      CCCA      C 3'
               CCUGCU      GGCC
               GGACGA      UCGG
miRNA  3' GACAA      CUUGAC    U 5'
```

```
position  1018
target 5'  C  GGA     C   A    G G 3'
            GU   UGCUG AGC GAGU A
            CA   ACGAC UUG CUCG U
miRNA  3' GA  AGG         A    G   5'
```

```
position  955
target 5'        A GG        G  C 3'
                  G  GGGCUGGG CG
                  C  CUUGACUC GU
miRNA  3' GACAAGGA GA        G    5'
```

```
position  344
target 5'    G     G     AUUCCCU    U 3'
              UCCUG  GAGC       GGCC
              AGGAC  CUUG       UCGG
miRNA  3' GACA     GA    AC         U 5'
```

```
position  141
target 5' U   C      A C  CCAAGG    A  3'
           CUG UCC GC G AC      GAGC
           GAC AGG CG C UG      CUCG
miRNA  3'     A   A  A U  A         GU 5'
```

**Suppl Fig 5: RNA hybrid microRNA target prediction algorithm identifies non-seed match miR-24-3p target sites in the 3'UTR of *Hmga1*.** We have identified eighteen non-seed match miR-24-3p target sites in the 3'UTR of *Hmga1.* Here, we have shown the top ten such miR-24-3p target sites.

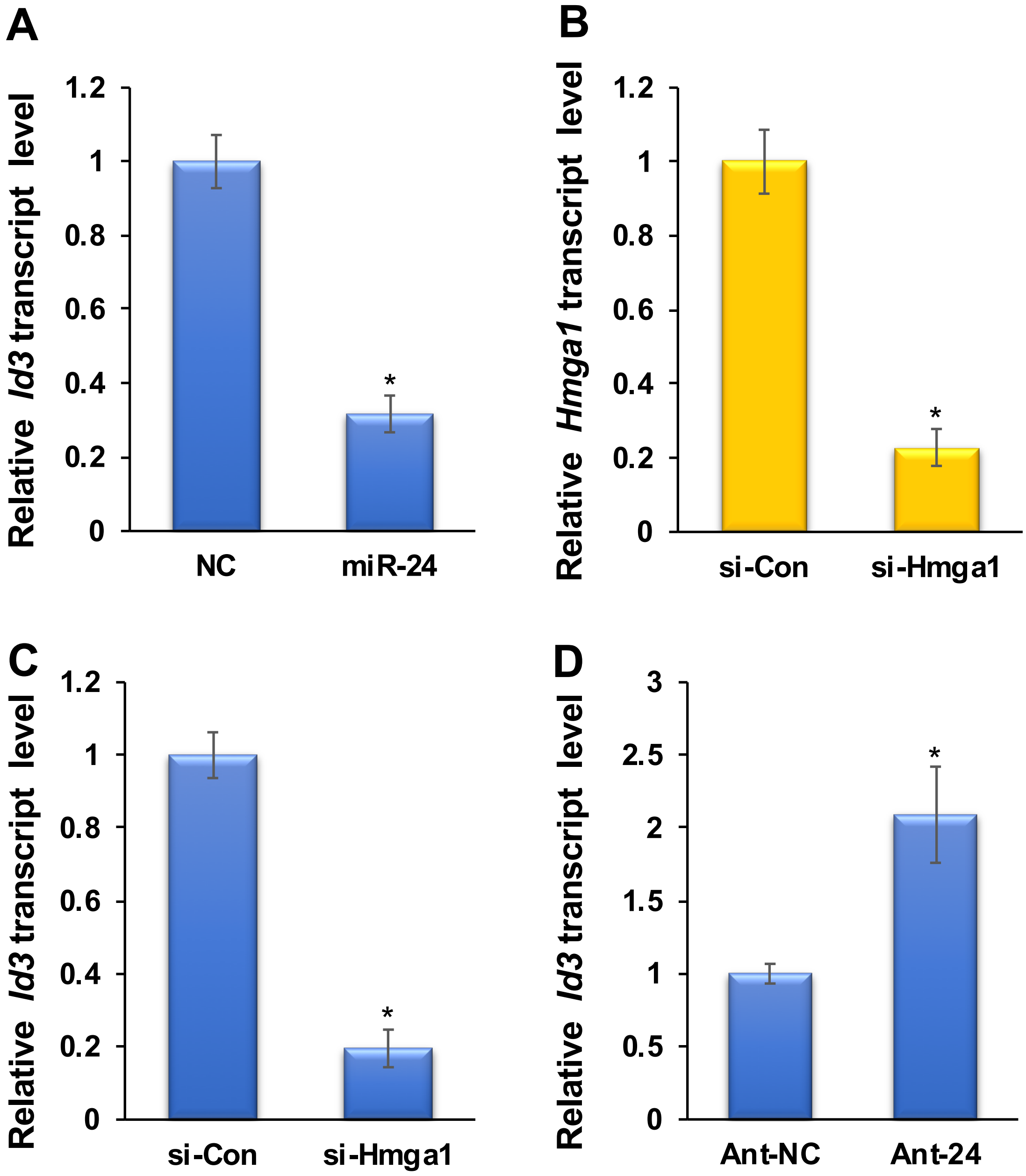


**Suppl Fig 6: miR-24-3p regulates HMGA1's direct downstream target *Id3* transcript level during myoblast differentiation.** (A) *Id3* transcript level is downregulated in miR-24-3p-transfected myoblasts. (B) *Hmga1* siRNA decreases *Hmga1* transcript level, and subsequently (C) decreases *Id3* transcript level. (D) Ant-24 derepresses *Id3* transcript level during myoblast differentiation. qRT-PCR was carried out for *Id3* or *Hmga1* and the values were normalized to the respective *Gapdh* values, then again to the values of the NC samples. The values are expressed as Mean ± SD of biological triplicates. *P < 0.001.

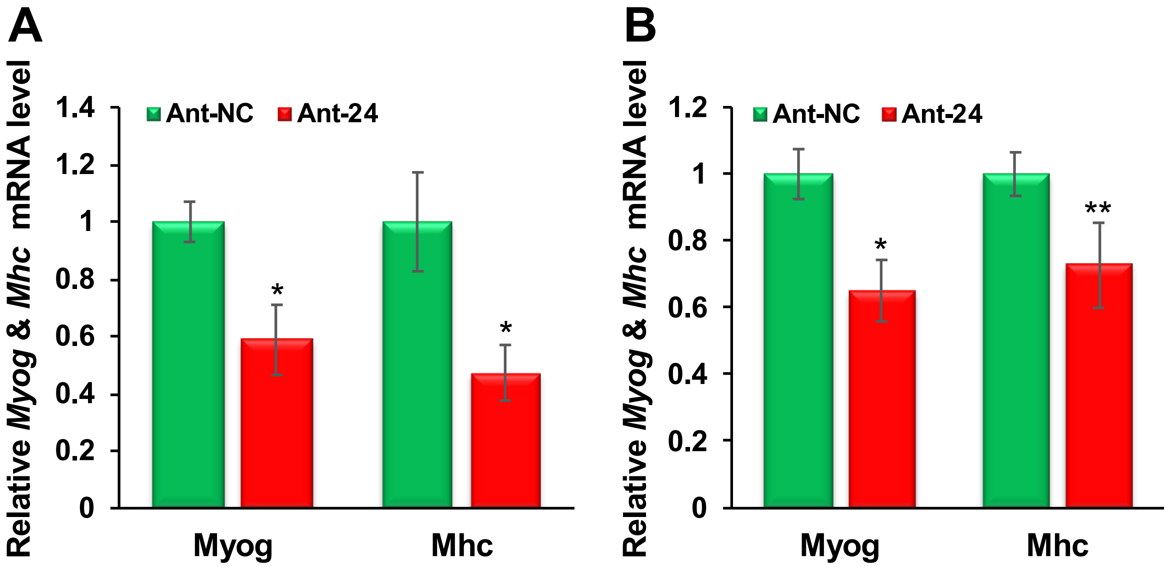


**Suppl Fig 7: *Myog* and *Mhc* levels are downregulated in the undifferentiated and differentiated MuSCs isolated from Ant-24-injected neonatal mice.** (A) *Myog* and *Mhc* mRNA levels are downregulated in the (A) undifferentiated and (B) differentiated (in DM for 48 hours) MuSCs. *Myog* and *Mhc* values were normalized with the respective RSP13 values, then again to the values of the Ant-NC samples. The values are expressed as Mean ± SD of five neonatal mice. *P < 0.001; **P<.0.01.

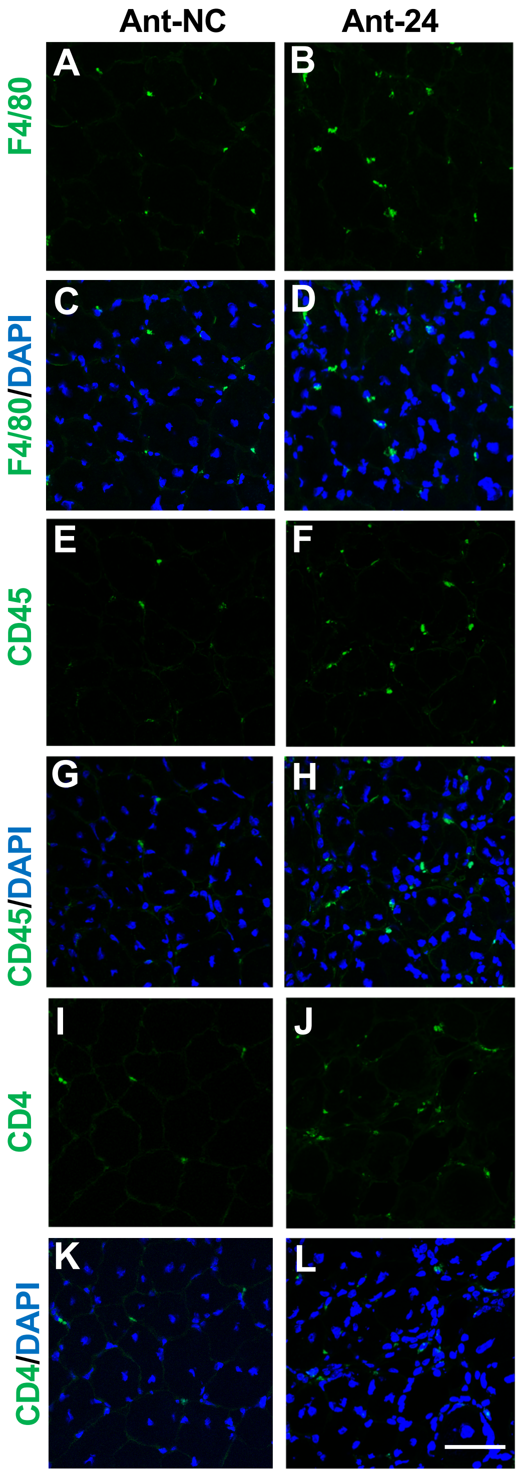


**Suppl Fig 8: Increased presence of various cells, including inflammatory cells are observed in the Ant-24-injected regenerating skeletal muscle.** Representative images of Ant-NC- and Ant-24-injected TA muscle sections immunostained with (A, B) F4/80, (C, D) F4/80 and DAPI, (E, F) CD45, (G, H) CD45 and DAPI, (I, J) CD4, (K, L) CD4 and DAPI . (A, C, E, G, I, K) Anti-NC-injected and (B, D, F, H, J, L) Ant-24-injected TA muscle sections on day 14 post-injury. Scale Bar: 50 μM.

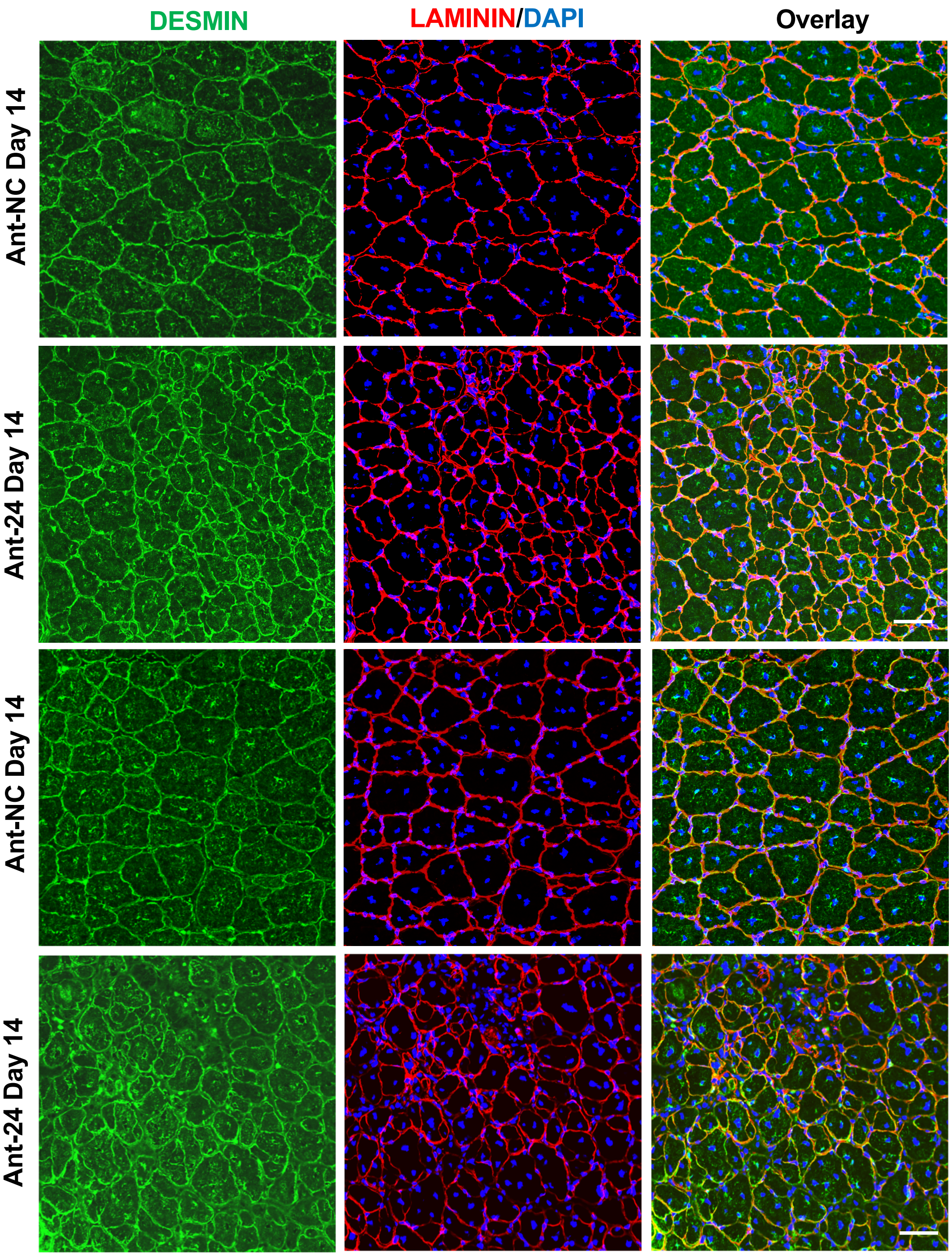


**Suppl Fig 9: Ant-24 impairs skeletal muscle regeneration after injury.** Additional representative images of Ant-NC- and Ant-24-injected TA muscle sections stained with DESMIN and LAMININ on day 14 post-injury. Ant-NC and Ant-24-injected samples are indicated. DESMIN (Green), LAMININ (red) and DAPI (Blue). Scale Bar: 50 μM.
